# The OSUMMER lines: a series of ultraviolet-accelerated NRAS-mutant mouse melanoma cell lines syngeneic to C57BL/6

**DOI:** 10.1101/2022.12.09.519766

**Authors:** Brandon M. Murphy, Daelin M. Jensen, Tiffany E. Arnold, Renan Aguilar-Valenzuela, Jase Hughes, Valentina Posada, Kimberly T. Nguyen, Vi T. Chu, Kenneth Y. Tsai, Craig J. Burd, Christin E. Burd

## Abstract

An increasing number of cancer subtypes are treated with front-line immunotherapy. However, approaches to overcome primary and acquired resistance remain limited. Pre-clinical mouse models are often used to investigate resistance mechanisms, novel drug combinations, and delivery methods; yet most of these models lack the genetic diversity and mutational patterns observed in human tumors. Here we describe a series of thirteen C57BL/6J melanoma cell lines to address this gap in the field. The Ohio State University-Moffitt Melanoma Exposed to Radiation (OSUMMER) cell lines are derived from mice expressing endogenous, melanocyte-specific, and clinically relevant *Nras* driver mutations (Q61R, Q61K, or Q61L). Exposure of these animals to a single, non-burning dose of ultraviolet B accelerates the onset of spontaneous melanomas with mutational patterns akin to human disease. Furthermore, in vivo irradiation selects against potent tumor antigens, which could prevent the outgrowth of syngeneic cell transfers. Each OSUMMER cell line possesses distinct in vitro growth properties, trametinib sensitivity, mutational signatures, and predicted antigenicity. Analysis of OSUMMER allografts shows a correlation between strong, predicted antigenicity and poor tumor outgrowth. These data suggest that the OSUMMER lines will be a valuable tool for modeling the heterogeneous responses of human melanomas to targeted and immune-based therapies.

**SIGNIFICANCE:** *NRAS*-activating mutations are the second most common genetic driver event in cutaneous melanoma, occurring in 15% to 25% of cases. With few therapeutic options beyond immunotherapy, patients with NRAS-mutant melanoma have a poorer prognosis. Pre-clinical mouse models that mimic the high mutational burden of human NRAS-mutant melanomas are lacking in the field. Here, we describe a series of NRAS-mutant melanoma cell lines, derived from ultraviolet (UV)-induced, spontaneous tumors. These lines permit the study of targeted, NRAS mutant-specific, immune, and combination therapies in C57BL/6J mice. With the release of this resource, we hope to catalyze new therapeutic approaches for NRAS-mutant melanoma.

## INTRODUCTION

Murine tumor cell lines play a vital role in advancing translational cancer research by providing a means to test the impact of genetic or pharmaceutical interventions both in vitro and in vivo. In particularly high demand are syngeneic tumor cell lines that can be used to test immune-based therapies which have become the standard of care for many solid tumors, including melanoma. Mouse B16 melanoma cells are a popular syngeneic cell line for pre-clinical immunotherapy studies. However, typical drivers of cutaneous melanoma (NRAS, BRAF, c-KIT) are not found in the B16 line, and in vivo expansion of these cells has been linked to viral components (Li et al., 1999; Mangeney et al., 2005). The availability of additional syngeneic lines reflective of human melanoma drivers and mutational patterns would facilitate relevant, robust, and reproducible pre-clinical studies.

The Yale University Mouse Melanoma (YUMM) series of cell lines began to address this gap in the field (Meeth et al., 2016). YUMM lines are derived from a series of genetically modified mouse models (GEMMs) carrying distinct combinations of cancer-associated alleles, including oncogenic *Braf*, *Pten* loss, *Cdkn2a* loss, or *Trp53* loss. Each YUMM cell line is syngeneic to C57BL/6J, allowing for the generation of tumors via subcutaneous injection into immune-competent hosts. One drawback of YUMM lines is that, unlike human melanomas, these cells have a low burden of single nucleotide variants (SNVs) (Alexandrov et al., 2013). To address this shortcoming, select YUMM cell lines were subjected to ultraviolet (UV) irradiation in vitro, yielding derivatives with a significant SNV burden and enhanced immunogenicity, termed the YUMM Exposed to Radiation or YUMM.ER lines (Wang et al., 2017). However, in the absence of a selection mechanism against highly immunogenic neoantigens, these cell lines may stimulate super-physiologic immune responses. Following the development of the YUMM.ER lines, Pérez-Guijarro et al., generated four syngeneic cell lines (C1-C4) from a series of *Braf-* and *Cdk4*-mutant mouse melanoma models irradiated in vivo (Pérez-Guijarro et al., 2020). Highlighting the potential power of such resources to advance clinical care, they defined a novel gene signature predictive of patient response to immune checkpoint blockade.

Unfortunately, syngeneic models representing BRAF-wild type melanoma remain limited. Of particular interest are models of NRAS-mutant melanoma, which constitutes between 15-25% of cases (Cancer Genome Atlas, 2015). Immunotherapy is the most common front-line treatment for these tumors because effective targeted therapies remain elusive (Johnson & Puzanov, 2015). Until recently, most NRAS-mutant mouse melanoma cell lines were derived from transgenic models that express the oncogene at super physiologic levels (Ackermann et al., 2005; Dorard et al., 2017; Petit et al., 2019; Swoboda et al., 2021). Such elevated expression alters RAS localization and signaling and may further sensitize the cells to pharmaceutical inhibitors (Prior & Hancock, 2012). Efforts by Bok et al. addressed this issue by establishing over 30 C57BL/6J melanoma cell lines expressing endogenous levels of oncogenic NRAS or BRAF (Bok et al., 2022; Bok et al., 2020). Notably, these cell lines contain a Cre-inducible reverse tetracycline-controlled transactivator (rtTA) and an integrated targeting site for recombination-mediated cassette exchange (RMCE), which make it possible to evaluate the impact of secondary genetic events in rapid succession. One drawback of these lines, however, is that they lack the high mutational burden and UV signature mutations observed in most human cutaneous melanomas. Given the frequent association of tumor mutational burden with immune checkpoint inhibitor responses (Klempner et al., 2020), cell lines that model the mutational landscape of human melanoma may be preferable for pre-clinical immunotherapy testing.

Here we describe the Ohio State University-Moffitt Melanoma Exposed to Radiation (OSUMMER) series of cell lines. OSUMMER cells are derived from spontaneous melanomas that arise in Tyr::Cre-ER(T2) *LSL-Nras* (*TN*) mice expressing one of four distinct NRAS mutants: Q61R, Q61K, Q61L, or G13R, from the endogenous gene locus. Subjecting these mice to a single, neonatal dose of UV irradiation leads to the formation of tumors with mutational patterns analogous to those observed in human melanoma (Bowman et al., 2021). Through in vitro, genomic, and in vivo studies, we show that OSUMMER cells have diverse growth rates, mutational burdens, MEK inhibitor sensitivities, and predicted antigenicity. These properties make the OSUMMER series well-suited for experiments to improve therapeutic options for NRAS mutant melanoma and other RAS-mutant tumor types.

## METHODS

### Mouse models

Animal research was conducted in accordance with protocols approved by the Ohio State Institutional Animal Care and Use Committee. The *TN^61R^* mouse model is homozygous for Tyr-CRE-ER(T2) (Bosenberg et al., 2006) and *LSL-Nras^61R^* (Burd et al., 2014) and was backcrossed seven generations to C57BL/6J mice. All other *TN* models were created using CRISPR-Cas9 to modify the endogenous *LSL-Nras* allele in *TN^61R^* mice as described (Murphy et al., 2022). CRISPR-modified mice were backcrossed two generations to the founding *TN^61R^* line to minimize off-target events. *Nras* expression was induced on postnatal days one and two via topical application of 20 mM 4-hydroxytamoxifen (4-OHT). On postnatal day three, the mice received one, 4.5 kJ/m^2^ dose of ultraviolet B (UVB) irradiation from a 16W, 312nm, fixed-position light source (see source details in: (Bowman et al., 2021)).

### Cell line derivation

Mice were euthanized and their melanomas resected when the tumors reached 0.8-1.6 cm^2^ in volume (length x width). A sample of approximately 5 mm^2^ was excised from each tumor and temporarily placed in a 15 ml conical tube containing 5 mL of ice-cold DMEM with 5% FBS, 1% penicillin-streptomycin, 1% L-glutamine and 1X antibiotic/antimycotic solution (Sigma A5955). Samples were then transferred to a 6-well plate containing 3 mL of PBS plus 1X antibiotic/antimycotic solution. Tumor samples were individually minced using 3-4 passes of a McIllwain tissue chopper (Ted Pella 10180) set to chop 150 mm sections at 60% of the maximal chopping rate and 50% force. Homogenized tumors were individually resuspended in 4 mL of digestion buffer (RPMI 1640, 10% FBS, 1% penicillin-streptomycin, 1% L-glutamine, 10 mg/mL collagenase type A, 0.25% trypsin, 0.02 mg/mL DNase I) and incubated in a 37°C water bath for 15 minutes. During the incubation period, the tube was inverted every 3-5 minutes. Dissociated cells were pelleted and then resuspended in melanocyte media (Hams F12 medium, 10% FBS, 7% horse serum, 1% pen-strep, 1% L-glutamine, 0.5 mM DB cAMP, 20 nM TPA, and 200 pM cholera toxin). The cell suspension was placed into a collagen-coated 10 cm dish. Once established (approximately 2-3 weeks), tumor cell lines no longer required collagen substrate and were maintained in DMEM containing 10% FBS, 1% penicillin-streptomycin, and 1% L-glutamine.

### Cell line validation

Cells were treated with 1% plasmocin (InvivoGen) and confirmed mycoplasma free using a PCR detection kit (ABM G238). Recombination of the *LSL-Nras* allele was verified by PCR using the following primers: Primer 1: 5’AGACGCGGAGACTTGGCGAGC-3’ Primer 2: 5’-GCTGGATCGTCAAGGCGCTTTTCC-3’ Primer 3: 5’-GCAAGAGGCCCGGCAGTACCTA-3’. PCR cycling conditions were as follows: 95°C 10 minutes, 35 x [95°C 15 seconds, 62°C 15 seconds, 72°C 10 seconds], 72°C 5 minutes. This PCR produces a 371 base pair product in the absence of recombination. A 562 base pair product is generated upon recombination and excision of the *LSL* cassette. NRAS protein expression was confirmed by immunoblot using cell lysate from the tumor cell lines. Cell lines were lysed in RIPA (25mM Tris pH 7.4, 150mM NaCl, 1% IGEPAL, 0.1% SLS) supplemented with protease inhibitor cocktail (Sigma P8340). Equal protein concentrations were run on an SDS-PAGE gel and transferred to PVDF (Sigma IPFL00010). PVDF membranes were blocked in 5% milk-PBST and then probed with NRAS (1:250, Abcam ab77392) and β-Actin (CST 3700S) primary antibodies in 5% BSA-PBST. Secondary antibodies were diluted in 5% milk-PBST as follows: anti-goat (1:15,000. LI-COR 926-32214), anti-mouse (1:15,000, LI-COR 926-68070). Membranes were imaged on a LI-COR Odyssey CLx system and quantified using Image Studio software (LI-COR Biosciences).

### Immunofluorescence

Each OSUMMER cell line was seeded in a 12-well plate at a density of 75,000-125,000 cells per well. Cells were grown overnight in DMEM supplemented with 10% FBS and 1x Glutamax (Gibco 35050-061). The next day, the cultures were washed in PBS, fixed in 4% paraformaldehyde, permeabilized in PBS containing 0.1% Tween-20 and 0.5% BSA, and blocked in 1% BSA. These cells were then incubated in a PBS solution containing 0.1% Tween-20, 1.0% BSA, and Phalloidin-Atto 488 (Sigma-Aldrich, 49409) and primary antibody (Anti-MelanA, Invitrogen, PIPA599174; Anti-PMEL, Invitrogen, PIPA5101023) at a 1:100 concentration. Control staining was conducted using a rabbit IgG (Millipore, NI01) antibody diluted at 1:100. Secondary antibody (Donkey anti-rabbit IgG Cy3-conjugate, Millipore, AP182C) was applied at a 1:500 dilution and the cells were counterstained with DAPI (Sigma-Aldrich, D9542). Images were taken on a Leica DMI 4000 inverted epifluorescence microscope using a 40x objective.

### Proliferation curves and IC_50_ determination

Cultured cell lines were trypsinized and counted on a hemocytometer. Each cell line was plated in triplicate at a density of 1,000 cells per well in a 96-well plate. The following day, cells were placed in the Incucyte ZOOM incubator and four 10x images of each well were taken every twenty minutes until the cells reached confluency. Data exported from the Incucyte ZOOM software package included average cellular confluency for each time point throughout the experiment. Growth curves were generated by plotting average confluency over time for each cell line. IC_50_ curves were generated by tracking the confluency of cells treated with 0.5, 1, 2, 4, and 8 μM trametinib (Selleckchem, S2673) over 48 hours using the Incucyte. At 48 hours, the confluency of each trametinib-treated well was normalized to the confluency of the vehicle-treated control. IC_50_ values were determined using GraphPad Prism version 8.4.3.

### Whole exome sequencing (WES) and mutational analyses

Genomic DNA was isolated from each cell line using the Zymo Quick-DNA Miniprep Kit (Zymo D3024) and quantified by Qubit. DNA concentration and quality were verified on an Agilent Tapestation prior to library generation. Bar-coded sequencing libraries were generated using SureSelect v6 Mouse All Exon and SureSelect QXT Library Prep kits and then sequenced on an Illumina HiSeq 4000. Paired-end, 150 bp reads were aligned to the mm39 mouse genome using the Burrows-Wheeler Aligner (BWA v0.7.17-r1188; (Li & Durbin, 2010)). Variants were called against pooled genomic DNA from *TN^61R^* breeders using Varscan2 (v2.4.3; (Koboldt et al., 2012)). Known polymorphisms were removed from the variant list. A list of potentially impactful variants was generated with Variant Effect Predictor (VEP; (McLaren et al., 2016)). Identified variants predicted to have a “moderate” or “high” impact by VEP were graphed in an Oncoprint using the ComplexHeatmaps package (version 2.0.0) in R (Gu et al., 2016). Ingenuity Pathway Analysis (QIAGEN Inc.; (Kramer et al., 2014)) was used to identify mutationally enriched gene ontologies. Single Base Substitution (SBS) signatures were probed using the SigProfiler Matrix Generator (Bergstrom et al., 2019).

### RNA Sequencing

The Zymo Quick-RNA Miniprep Kit (Zymo R1054) was used to isolate RNA from each cell line. RNA concentration was quantified by Qubit and sample quality was verified on an Agilent Tapestation prior to library generation. Library preparation and RNA sequencing was conducted by Novogene Co. (Novogene Co., USA) on the Illumina NovaSeq 6000 platform. Paired-end, 150 bp reads were aligned to the mm39 mouse genome using STAR (v2.7.9a; (Dobin et al., 2013)). Gene expression and coverage were assessed using Cufflinks (Trapnell et al., 2012). Gene expression patterns enriched in trametinib-resistant OSUMMER cell lines were identified using the ShinyGo tool (Ge et al., 2020).

### Neoantigen Prediction

Neoantigen prediction was performed according to the pVACseq workflow (Hundal et al., 2016); https://github.com/CEBurdLab/OSUMMERNeoantigenPrediction). First, the list of VEP-annotated variants identified by WES were annotated with RNA-seq expression data using vatools (www.vatools.org). Next, potential neoantigens were predicted in silico based on binding affinity to H2-K^b^, H2-D^b^, and H2-IA^b^ (Immune Epitope Database; (Vita et al., 2019)). Prediction algorithms included MHCflurry (O’Donnell et al., 2020), MHCnuggets (Shao et al., 2020), NNalign (Nielsen & Andreatta, 2017), NetMHC (Lundegaard et al., 2008), SMM, SMMPMBEC, and SMMalign (Nielsen et al., 2007). Neoepitopes were identified and filtered based on the median MHC binding affinity predicted across all seven algorithms, as well as DNA and RNA read depth, and mRNA expression data.

### OSUMMER Tumor Allografts

Cell lines were trypsinized, pelleted, and washed twice with 1x PBS. Cells were then counted on a hemocytometer and resuspended in 1x PBS at a concentration of 200,000 cells per mL. 100 μL of the resulting cell solution was injected into each flank of anesthetized C57BL/6J mice. In each case, the sex of the C57BL/6J recipients matched that of the injected cell line. Mice were monitored every other day until tumor formation. Once developed, tumors were measured daily using digital calipers. Tumor size was calculated as length x width. Mice were euthanized upon reaching predetermined exclusion criteria, and the tumors were excised for downstream analyses.

### Flow Cytometry

Excised tumors were mechanically chopped with surgical scissors, and then enzymatically dissociated in PBS containing 0.25% Collagenase D (Millipore Sigma, 11088866001) and 10,000 units of DNase (Omega BioTek, #A122111QGPCSDA03) per milliliter. Cells were passed through a 70 μm cell strainer to filter out any remaining chunks of tissue. Immune cells were isolated using a Ficoll gradient as described by the manufacturer (GE Healthcare, GE17-0891-01). The resulting immune cells were resuspended in 1x PBS prior to being plated in a 96-well plate for flow staining.

Immune cells were first incubated with Fc block (anti-CD16/CD32 1:100; BD Biosciences #553142) and then stained with one of two immunophenotyping panels diluted in 1x FACS buffer (2% FBS, 0.2% sodium azide in PBS). The lymphoid panel consisted of antibodies directed against PD-1 (Biolegend #135241; 1:250), FoxP3 (Biolegend #126419; 1:250), Lag-3 (Biolegend #C9B7W; 1:250), CD3 (BD Biosciences #145-2C11; 1:250), CD4 (BD Biosciences #563933; 1:250), CD45 (Tonbo Biosciences #50-0451; 1:250), and CD8 (Tonbo Biosciences #20-1886; 1:250) as well as GhostDye viability dye (Tonbo Biosciences #13-0865; 1:1000). The myeloid panel contained GhostDye (1:1000) and antibodies directed against CD45 (Tonbo Biosciences #50-0451; 1:250), CD11b (Tonbo Biosciences #75-0112; 1:250), CD11c (Tonbo Biosciences #60-0114; 1:250), F4/80 (Tonbo Biosciences #20-4801; 1:250), Ly6G (Tonbo Biosciences #25-1276; 1:250), and Ly6C (Biolegend #128049; 1:250). Single color controls were made using UltraComp eBeads (Invitrogen # 01-2222-41). Data were acquired on a BD Fortessa II flow cytometer using BD FACSDiva software (BD Biosciences) and analyzed using FlowJo software (Tree Star).

## RESULTS

We used the *TN^61R^, TN^61L^, TN^61K^*, and *TN^13R^* mouse models to generate NRAS-mutant melanoma cell lines syngeneic to C57BL/6J (Murphy et al., 2022). Each mouse model is homozygous for a melanocyte-specific Cre-ER(T2) transgene (Tyr::Cre-ER(T2); (Bosenberg et al., 2006)) and a conditional *LSL-Nras* allele encoding either NRAS Q61R, Q61K, Q61L, or G13R (**Figure 1A**). Notably, NRAS Q61R, Q61K, and Q61L mutants occur frequently in human melanoma, whereas NRAS G13R mutants are rare (**Figure 1B**). Experimental *TN* pups were treated with 4-hydroxytamoxifen (4OHT) on postnatal days one and two to activate Cre recombinase and induce mutant NRAS expression in melanocytes (**Figure 1C**). On postnatal day three, the mice were subjected to a single, non-burning dose of UVB which mimics the etiologic role of sunlight in melanoma. Cell lines were generated from spontaneous melanomas that arose 5 to 32 weeks after irradiation (**Figure 1D**). The melanocytic origin of each cell line was verified by assessing NRAS protein expression, *LSL* recombination, and MelanA or PMEL staining (**Figure 1E-F**; **Supplemental Figure S1**). All thirteen cell lines adopted a dendritic phenotype typical of cultured melanocytes (**Supplemental Figure S2**). **Table 1** summarizes the sex, genotype, and genomic characteristics of each OSUMMER line.

**Figure 1.**
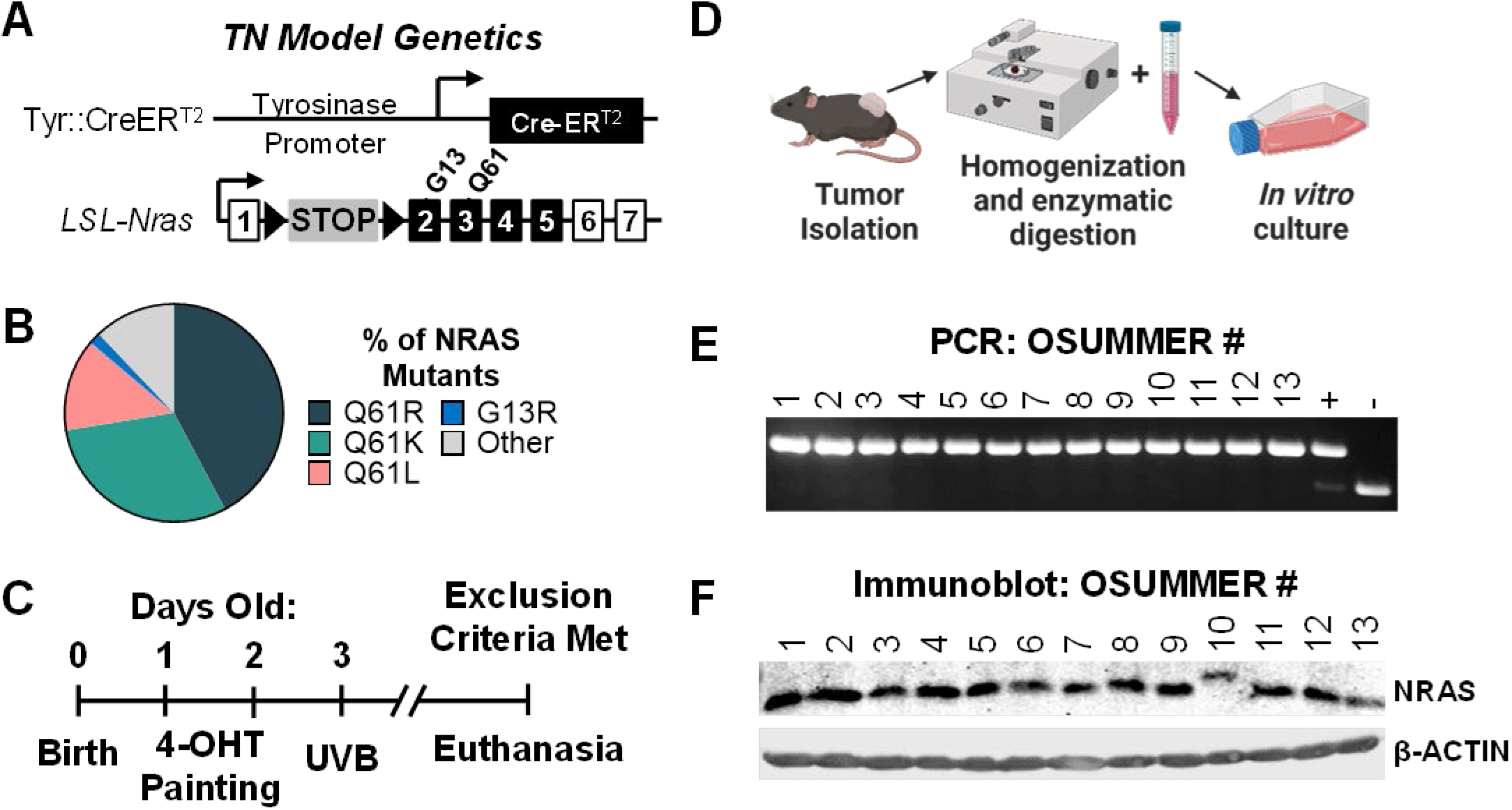
Generation and validation of the OSUMMER cell lines. **A,** Diagram of the homozygous alleles present in the *TN* mouse models. Tyr::CreER^T2^ drives the expression of a tamoxifen-inducible Cre recombinase specific to melanocytes. A floxed transcriptional and translational stop cassette (*LSL*) is inserted into the endogenous *Nras* promoter to prevent gene expression in the absence of Cre. *Nras* mutations encoding Q61R, Q61K, Q61L, or G13R were engineered into the endogenous gene locus of each *LSL-Nras* allele. **B,** Frequency of specific amino acid substitutions among NRAS-mutant, human cutaneous melanomas in the National Cancer Institute’s Genomic Data Commons Data Portal (n = 116 tumors; (Grossman et al., 2016)). **C,** Experimental setup to generate spontaneous NRAS-mutant melanomas. *TN* mice are painted with 4-hydroxytamoxifen (4-OHT) on postnatal days one and two to induce Cre activity, *LSL* recombination, and mutant NRAS expression. On postnatal day three, the animals receive a single, non-burning dose of ultraviolet B (UVB) irradiation. Animals are monitored for spontaneous melanoma formation and tumors are harvested for cell line generation before exceeding the predetermined exclusion criteria. **D,** Diagram of tumor processing for OSUMMER cell line generation. Additional details can be found in the Methods. **E,** PCR screen to confirm *LSL* recombination in genomic DNA isolated from the OSUMMER cell lines. The expected product sizes for the native and recombined *LSL-Nras* alleles are 371 and 562 bp, respectively. + = Homozygous *LSL-Nras* mouse embryonic fibroblasts (MEFs) treated with adenoviral Cre. - = *LSL-Nras* MEFs prior to recombination. **F,** Immunoblot of protein lysates isolated from the OSUMMER cell lines. Note that melanoma-associated NRAS codon 61 mutants are known to migrate faster than codon 13 mutants during SDS-PAGE (Der et al., 1986; Murphy et al., 2022).

**Table 1.**
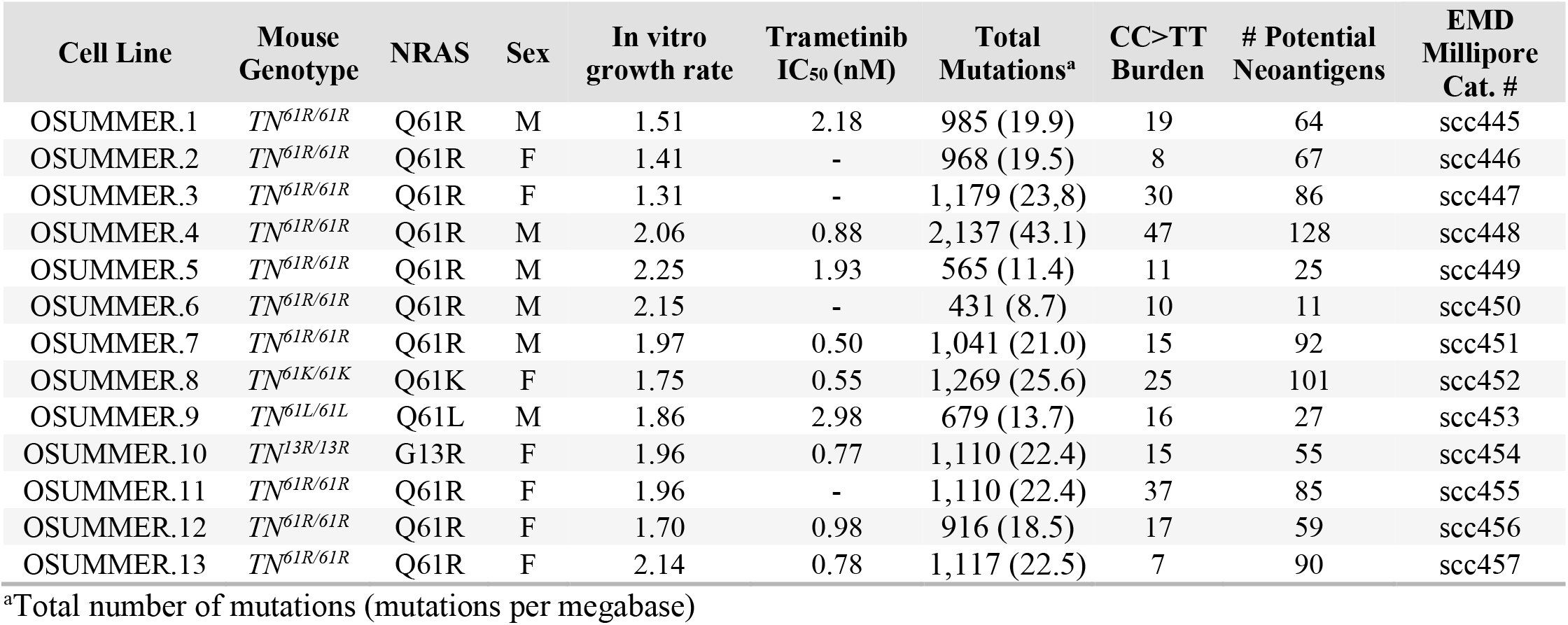
Summary of OSUMMER cell line characteristics.

| Cell Line | Mouse Genotype | NRAS | Sex | In vitro growth rate | Trametinib IC<sub>50</sub> (nM) | Total Mutations<sup>a</sup> | CC>TT Burden | # Potential Neoantigens | EMD Millipore Cat. # |
| --- | --- | --- | --- | --- | --- | --- | --- | --- | --- |
| OSUMMER.1 | *TN<sup>61R/61R</sup>* | Q61R | M | 1.51 | 2.18 | 985 (19.9) | 19 | 64 | scc445 |
| OSUMMER.2 | *TN<sup>61R/61R</sup>* | Q61R | F | 1.41 | - | 968 (19.5) | 8 | 67 | scc446 |
| OSUMMER.3 | *TN<sup>61R/61R</sup>* | Q61R | F | 1.31 | - | 1,179 (23.8) | 30 | 86 | scc447 |
| OSUMMER.4 | *TN<sup>61R/61R</sup>* | Q61R | M | 2.06 | 0.88 | 2,137 (43.1) | 47 | 128 | scc448 |
| OSUMMER.5 | *TN<sup>61R/61R</sup>* | Q61R | M | 2.25 | 1.93 | 565 (11.4) | 11 | 25 | scc449 |
| OSUMMER.6 | *TN<sup>61R/61R</sup>* | Q61R | M | 2.15 | - | 431 (8.7) | 10 | 11 | scc450 |
| OSUMMER.7 | *TN<sup>61R/61R</sup>* | Q61R | M | 1.97 | 0.50 | 1,041 (21.0) | 15 | 92 | scc451 |
| OSUMMER.8 | *TN<sup>61K/61K</sup>* | Q61K | F | 1.75 | 0.55 | 1,269 (25.6) | 25 | 101 | scc452 |
| OSUMMER.9 | *TN<sup>61L/61L</sup>* | Q61L | M | 1.86 | 2.98 | 679 (13.7) | 16 | 27 | scc453 |
| OSUMMER.10 | *TN<sup>13R/13R</sup>* | G13R | F | 1.96 | 0.77 | 1,110 (22.4) | 15 | 55 | scc454 |
| OSUMMER.11 | *TN<sup>61R/61R</sup>* | Q61R | F | 1.96 | - | 1,110 (22.4) | 37 | 85 | scc455 |
| OSUMMER.12 | *TN<sup>61R/61R</sup>* | Q61R | F | 1.70 | 0.98 | 916 (18.5) | 17 | 59 | scc456 |
| OSUMMER.13 | *TN<sup>61R/61R</sup>* | Q61R | F | 2.14 | 0.78 | 1,117 (22.5) | 7 | 90 | scc457 |

<sup>a</sup>Total number of mutations (mutations per megabase)

Mitogen-activated protein kinase (MAPK) signaling is a hallmark of melanoma development and a frequent target of pharmaceutical inhibitors (Downward, 2003; Shain et al., 2018). However, the success of agents like the Mitogen-activated protein kinase kinase (MEK) inhibitor, trametinib (GSK1120212), in preclinical studies has not translated into clinical benefit for patients with NRAS mutant melanoma (Dummer et al., 2017; Falchook et al., 2012; Gilmartin et al., 2011; Kwong et al., 2012; Lebbe et al., 2020; Zimmer et al., 2014). To test whether a subset of OSUMMER cell lines might recapitulate clinical experiences with trametinib, we evaluated the trametinib sensitivity of nine OSUMMER lines using Incucyte live-cell imaging. Trametinib IC_50_ values varied from 0.5 to 2.9 nM and were independent of culture growth rate or the underlying *Nras* mutation (**Table 1**). We next performed RNA-Seq on all 13 OSUMMER cell lines to identify gene expression signatures associated with trametinib resistance. 312 transcripts differed by 1.5-fold (padj < 0.001) between trametinib-sensitive and -resistant (IC_50_ > 1.9 nM) OSUMMER cell lines (**Supplemental Table 1**). Pathways upregulated in trametinib-sensitive cell lines included those associated with neuronal development/differentiation and cytokine responses (**Supplemental Table 1**). Transcripts associated with trametinib resistance included *Hoxd8* and *Hoxd9* (up 8.8- and 12.1-fold, respectively in resistant lines; **Supplemental Table 1**). Prior publications have identified melanocyte plasticity and HOXD8 overexpression as mechanisms of MAPK pathway inhibitor resistance (Olbryt et al., 2020; Van Allen et al., 2014; Whittaker et al., 2013). Our data support these observations and suggest that the heterogeneity of the OSUMMER cell lines will be advantageous for studying drug sensitivity and resistance mechanisms.

Knowledge of the secondary mutations present in each OSUMMER cell line would aid in the design and interpretation of studies using this resource. Therefore, we performed whole-exome sequencing (WES) of all thirteen OSUMMER cell lines. A complete listing of mutations identified in the OSUMMER lines appears in **Supplemental Table 2**. Recurrent mutations among the cell lines included those located in large, repetitive genes and several genes identified in prior genomic analyses of *TN* tumors (**Figure 2**; (Bowman et al., 2021)). Specifically, 77% (10/13) of OSUMMER cell lines and 57% (4/7) of *TN* primary tumors contained *Flna* mutations predicted to impair F-Actin binding, mRNA translation, or protein stability (e.g., L1170P/R, S1197S/N, S1199F, I1219F; (Bowman et al., 2021)). Mutations in the RING domain of Mitogen-activated protein kinase kinase kinase 1 (MAP3K1) were also seen in both datasets. In contrast, *trp53* mutations were specific to the OSUMMER cell lines and not seen in our prior analyses of 17 UV-accelerated primary *TN* tumors (Bowman et al., 2021; Murphy et al., 2022). The *trp53* mutants found in OSUMMER.4, OSUMMER.7, and OSUMMER.8 were analogous to DNA binding domain mutations reported in human cancer (F134V, K120M, C135W; (de Andrade et al., 2022)). Thus, it appears that in vitro culture selects for the outgrowth of *TN* melanoma cells with impaired p53 function.

**Figure 2.**
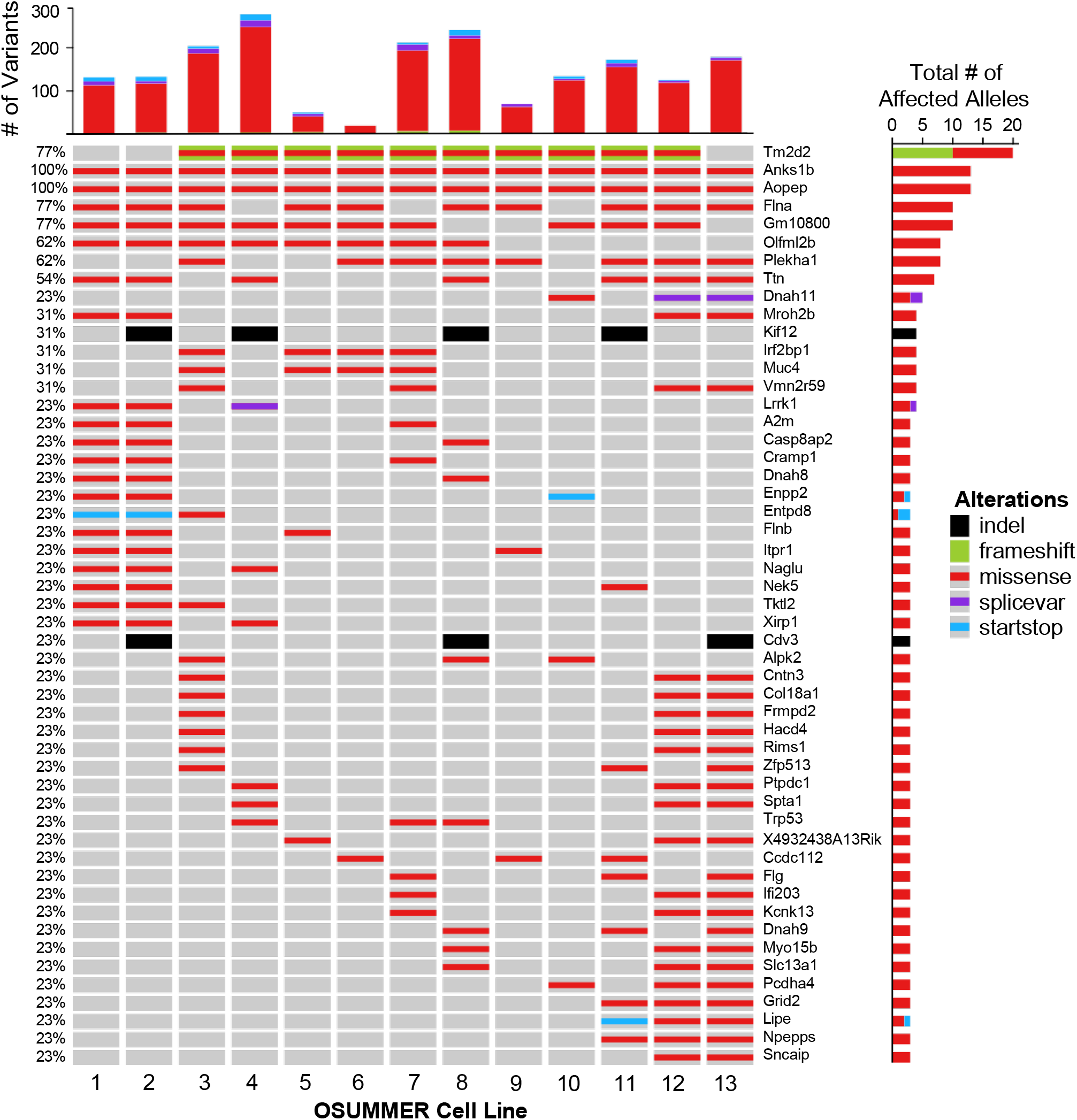
Recurrent genetic alterations in the OSUMMER cell lines. ***Top***, Bar graph depicting the total number of gene variants with moderate to high impact in each OSUMMER cell line. Gene variants were determined by WES and annotated using the Variant Effect Predictor as described in the Methods. ***Bottom,*** Oncoprint of genes mutated in at least two OSUMMER cell lines. Colors indicate the mutation type: missense (red), splice variant (purple), or nonsense/stop (blue). The frequency and number of OSUMMER cell lines containing a variant in each listed gene are indicated on the left and right of each corresponding row.

Exposure to UV irradiation promotes the acquisition of C>T transition mutations. Accumulation of these and other DNA alterations leads to a high number of single nucleotide variants (SNVs) in sun-exposed melanomas (Alexandrov et al., 2013). However, GEMM-derived spontaneous tumors are often characterized by genomic rearrangements rather than SNVs (Bowman et al., 2021; McFadden et al., 2016; Westcott et al., 2015). We previously reported that the mutational signatures present in UV-accelerated *TN* tumors are analogous to those observed in human melanoma (Bowman et al., 2021). Therefore, we wondered if the OSUMMER cell lines would retain these signatures in vitro. The total mutational burden of the OSUMMER cell lines ranged from 431 to 2,137 variants per cell line or 8.7 to 43.1 variants per megabase (**Figure 3A**; **Table 1**). For reference, the median total mutation burden of human skin melanoma is 14.4 variants per megabase (95% CI = 36.5-43 variants/Mb; (Chalmers et al., 2017)). Genomic evidence of prior UV exposure was more prominent in cell lines like OSUMMER.4 where C>T transition and dipyrimidine mutations predominated (**Figure 3B-C**; **Table 1**). Furthermore, the contribution of mutational patterns associated with UV carcinogenesis in humans, SBS7a and 7b (Alexandrov et al., 2013), also varied among the OSUMMER cell lines (**Figure 3D**). These data show a diversity in the extent to which OSUMMER cell lines retain evidence of prior UV exposure and mutational patterns analogous to those observed in human melanoma.

**Figure 3.**
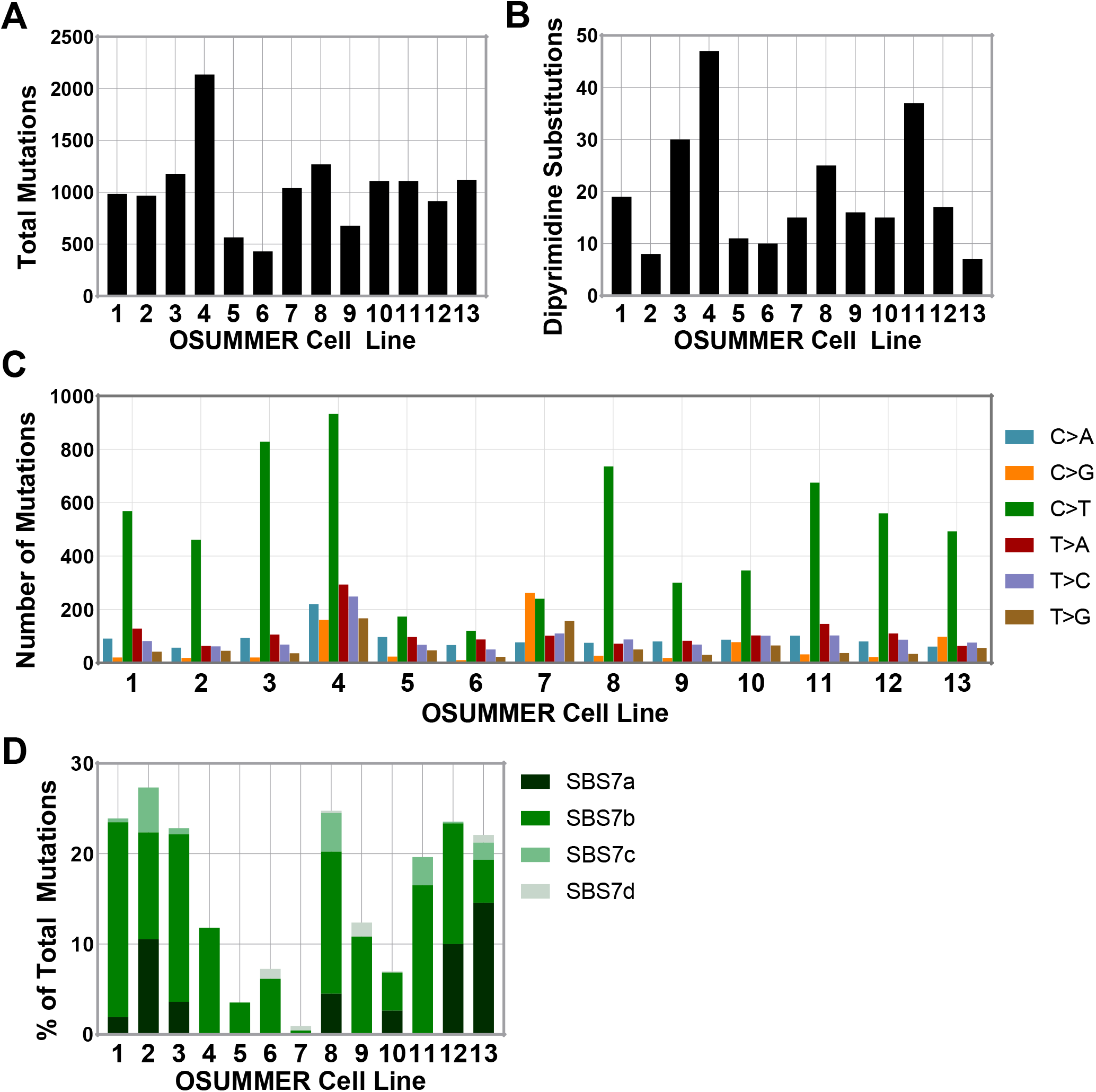
Total mutation burden and number of UV signature mutations vary among the OSUMMER cell lines. **A,** Total mutation burden as determined by WES of each OSUMMER cell line. **B,** Number of dipyrimidine mutations, indicative of faulty UV damage repair, detected by WES of each OSUMMER cell line. **C,** Distribution of mutation types detected in WES from each OSUMMER cell line. C>T transition mutations are often associated with UV damage, whereas C>A transversions are characteristic of oxidative damage. **D,** Percentage of mutations in each cell line explained by COSMIC mutational signatures associated with UV damage (i.e. SBS 7a-d).

Not all somatic mutations generate neoantigens. Therefore, we assessed the potential antigenicity of mutations in each OSUMMER cell line using the pVAC-Seq pipeline (Hundal et al., 2016; Hundal et al., 2020). pVAC-Seq combines somatic mutation calling, mRNA expression data, and major histocompatibility complex (MHC) epitope prediction to identify and prioritize putative neoantigens (see Methods). The percentage of neoantigenic variants predicted by pVAC-Seq varied among the OSUMMER cell lines (**Figure 4A**, **Supplemental Table 3**). Of the variants predicted to alter peptide sequence, 4.5 to 16.6% were identified as having the potential to generate neoantigens. The total number of MHC-binding neoantigen peptides was greatest in OSUMMER.4 (128 peptides) and OSUMMER. 8 (101 peptides) cells and lowest in OSUMMER.5 (25 peptides), OSUMMER.6 (11 peptides), and OSUMMER. 9 (27 peptides) cells (**Figure 4B**; **Table 1**). Neoantigen burden paralleled the mutational burden of each cell line, consistent with the idea that there is a greater likelihood of generating neoantigens in tumors with a high mutational load (**Figure 4C**).

**Figure 4.**
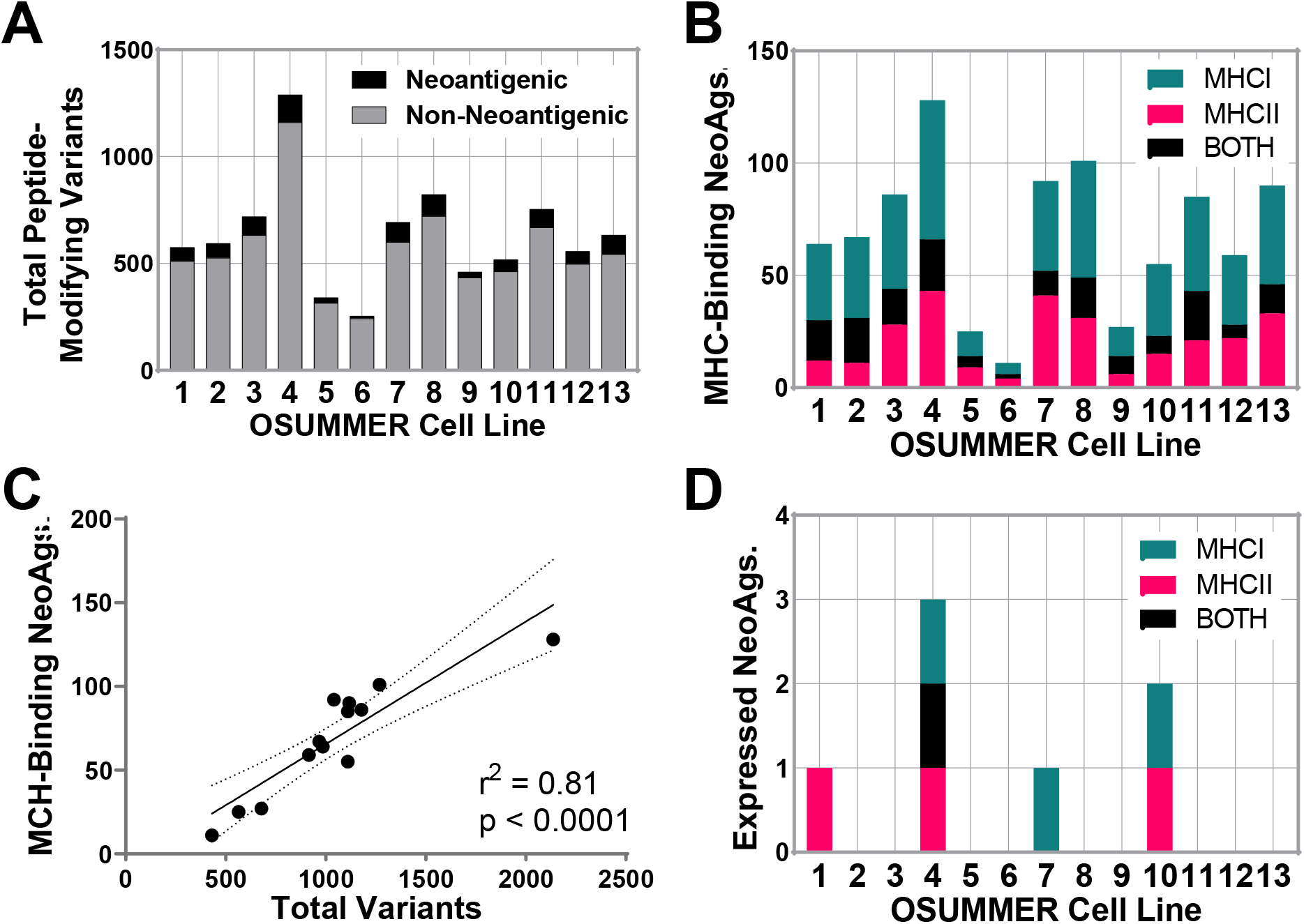
Predicted neoantigen burden differs among the OSUMMER cell lines. **A,** Graph of the total peptide-modifying genomic variants identified in each OSUMMER cell line using the Ensembl VEP. The neoantigenic potential of each modified peptide was determined using pVACseq. **B,** Graph of the total number of predicted neoantigens in each OSUMMER cell line classified by their predicted MHC (MHCI: H2-K^b^, H2-D^b^, MHCII: H2-IA^b^) binding affinity. **C,** Linear regression of the total number of variants plotted against the number of MHC-binding neoantigens in each OSUMMER cell line. **D,** Graph of the total number of expressed neoantigens as determined by incorporating RNAseq data from each OSUMMER cell line into the pVACseq algorithm.

We filtered the list of potential neoantigens in each OSUMMER cell line using RNA-Seq data to eliminate any variants that were not expressed in vitro. After filtering, OSUMMER.1, OSUMMER.4, OSUMMER.7, and OSUMMER.10 were the only lines with predicted MHC-binding neoantigens expressed in culture (**Figure 4D**). None of the expressed neoantigens overlapped between cell lines, and the number of expressed neoantigens did not correlate with the mutational burden of each cell line. However, we cannot rule out the possibility that an expanded repertoire of neoantigens would be observed in vivo or in proteomic analyses of MHC complexes. Together, these data highlight the diversity in putative neoantigenic burden among OSUMMER cell lines.

Tumors enriched in neoantigens should be more immunogenic. To examine whether this correlation held true in OSUMMER cells, we injected lines predicted to have high (OSUMMER.4), medium (OSUMMER.2, OSUMMER.3), and low (OSUMMER.5) antigenicity into syngeneic, C57BL/6J mice. Injected cell lines were resuspended in sterile saline to limit any immunological effects of substrates like Matrigel. Tumor growth rates partially paralleled the predicted antigenicity of each cell line with OSUMMER.5 growing the fastest and OSUMMER.4 growing the slowest (**Figure 5A**). More striking was the difference in tumor take rates, where the vast majority of OSUMMER.4 injections did not result in persistent tumor outgrowth (**Figure 5B**). These data support a general correlation between predicted antigenicity and tumor growth potential among the OSUMMER lines.

**Figure 5.**
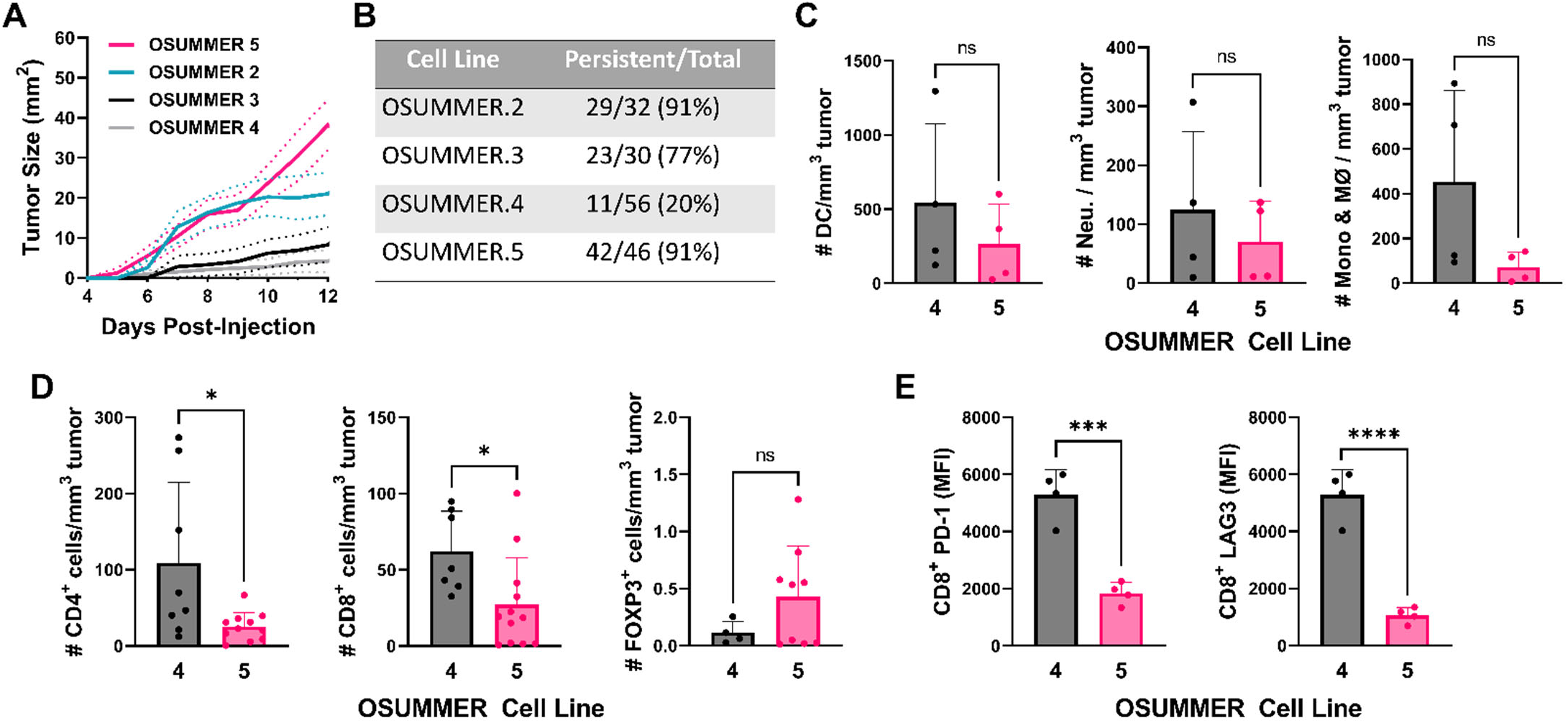
Tumor outgrowth and immune infiltration differ between OSUMMER cell lines. **A,** Growth curve of OSUMMER.2, OSUMMER.3, OSUMMER.4, and OSUMMER.5 tumors in C57BL/6J recipients. Filled lines represent the average growth rate of the persistent tumors indicated in ‘B’. Dotted lines show the 95% confidence intervals. **B,** Table showing the number of cell injections that resulted in persistent tumor growth (reaching exclusion criteria) for each OSUMMER cell line. **C,** Dendritic cell (DC, p = 0.389), neutrophil (Neu, p = 0.500), and monocyte and macrophage (Mono & MØ, p = 0.113) densities per mm^3^ of tumor as determined by flow cytometry. **D,** Density of CD4^+^ T cells (p = 0.247), CD8^+^ T cells (p = 0.6), and CD4^+^ FOXP3^+^ regulatory T cells (p = 0.458) per mm^3^ of tumor as assessed by flow cytometry. **E,** PD-1 (p = 0.0004) and LAG3 (p = <0.0001) positivity in CD8^+^ T cells expressed as median fluorescence intensity. **Statistics for C-E,** Each dot represents a distinct tumor. Bar heights indicate the data mean, and error bars show the standard deviation. Values for the OSUMMER.4 and OSUMMER.5 cell lines were compared using unpaired t-tests.

We wondered if tumor cell lines with high predicted antigenicity and low tumor take would elicit a more potent immune response. Therefore, we compared the immune infiltrates in OSUMMER.4 (high burden) and OSUMMER.5 (low burden) tumors. Dendritic cell, neutrophil, and monocyte/macrophage numbers did not differ significantly between OSUMMER.4 and OSUMMER.5 tumors (**Figure 5C**; **Supplemental Figure S3**). However, lymphocyte populations differed significantly between tumors generated from each cell line. OSUMMER.4 tumors had more CD4+ and CD8+ T cell infiltrates (**Figure 5D**). Infiltrating CD8+ T cells expressed higher levels of the immune checkpoint molecules, PD-1 and Lag-3, in OSUMMER.4 tumors (**Figure 5E**). These data, along with the limited outgrowth of injected OSUMMER.4 cells, suggest that tumor outgrowth may be dependent upon the establishment of an exhausted T cell phenotype. Overall, our allograft results highlight distinctions among the immune responsiveness of OSUMMER tumors which may aid in the testing of different immunotherapeutic modalities for NRAS mutant melanoma.

## DISCUSSION

The OSUMMER cell lines are a new resource for testing single-agent and combination therapies for NRAS-mutant melanoma in immune-competent models. The most common NRAS mutants in human melanoma, Q61R, Q61K, and Q61L, are represented in the OSUMMER series, along with the most frequently occurring codon 12/13 mutant, G13R (Grossman et al., 2016). This variety of driver alleles will benefit ongoing efforts to develop and test mutant-specific RAS inhibitors (Molina-Arcas et al., 2021). Expression of NRAS from the endogenous gene locus ensures that protein localization and signaling remain at physically relevant levels. Additional attributes of the OSUMMER lines, including knowledge of the secondary gene mutations and transcriptional profiles of each cell line, may help to identify primary drug resistance mechanisms.

Our prior work established that a single, neonatal dose of UV irradiation dramatically accelerated the formation of spontaneous melanomas with mutational patterns analogous to human disease (Bowman et al., 2021; Hennessey et al., 2017; Murphy et al., 2022). We speculated that these genomic patterns would be retained in the OSUMMER cell lines. Indeed, the total mutation burdens and UV-associated COSMIC mutational signatures found in OSUMMER cell lines mimic those of human cutaneous melanoma (**Figure 3**, **Table 1**). While similar genomic signatures are seen in the YUMM.ER lines, which were irradiated ex vivo (Wang et al., 2017), we believe that in vivo UV irradiation has several advantages. First, melanocytes that acquire strong neoantigens may be eliminated by the immune system prior to melanoma onset. This process could prevent the outgrowth of clones that elicit a potent immune reaction. Second, in vivo selection may enrich UV-induced mutations that promote tumorigenesis. Therefore, a higher percentage of mutations in the OSUMMER lines could play a role in tumor progression. In support of this idea, cell lines derived from UV-irradiated Tyr::Cre-ER(T2); *Braf*^*CA*/+^; *Pten*^*fl*/+^; *Cdkn2a*^*fl*/+^, Tyr::Cre-ER(T2); Hgf+; *Braf*^*CA*/+^; *Cdkn2a*^*fl*/+^, and Hgf+ mice yielded tumors with mutational profiles similar to the major genetic subtypes of human melanoma, BRAF-mutant, RAS-mutant, and triple wildtype (Pérez-Guijarro et al., 2020). Together, these data highlight how *in vivo* UV exposure may distinguish the OSUMMER series from other NRAS-mutant melanoma cell lines.

Immunotherapy is the standard of care, first-line therapy for NRAS-mutant melanoma. However, few second-line options are available for these patients, and well-characterized pre-clinical models for immunotherapeutic testing are limited. While UV-accelerated melanoma GEMMs can be used for immunotherapeutic studies, tumors in these models arise asynchronously, in random locations, and grow at different rates. With the idea that the OSUMMER lines would be used to develop and test immune, targeted, and combination therapies, we sought to characterize their potential immunogenicity using the pVAC-Seq pipeline (Hundal et al., 2016; Hundal et al., 2020). We found that the frequency of potential neoantigenic peptides varied among the OSUMMER cell lines, ranging from 4.3-14.2% of the nonsynonymous variants (SNVs, indels, splice site mutations, gene fusions) detected in each culture (**Figure 4A**). Even with this variance, the number of predicted neoantigens directly paralleled the mutational burden of each OSUMMER cell line (**Figure 4C**). These data were consistent with the use of tumor mutational burden to predict response to immune checkpoint inhibitor therapy (Strickler et al., 2021).

Although the number of predicted neoantigens paralleled the mutational burden of each OSUMMER cell line, most of these antigen RNAs were not expressed in vitro (**Figure 4B, D**). These data highlight a potential disconnect between tumor mutational burden and antigenicity. However, transcript abundance may not be the most accurate predictor of antigenic potential. A recent study by Jaeger et al. used purified MHCI complexes from autochthonous mouse pancreas and lung adenocarcinomas to show that neither mRNA level, nor translational efficiency is a reliable predictor of antigen presentation in vivo (Jaeger et al., 2022). In fact, many of the presented peptides were rare transcripts expressed at low levels, suggesting a role for post-translational mechanisms in shaping tumor-specific neoantigen presentation. Our tumor growth data support these findings, showing that the growth of OSUMMER.5 and OSUMMER.2, differs drastically from OSUMMER.3 even though all three cell lines are predicted to express no neoantigens (**Figures 4D**, **5A**). Another possibility is that environmental cues may alter the expression of variant alleles in vitro and in vivo. Therefore, it may be worthwhile to consider all possible antigenic variants when analyzing data from the OSUMMER lines, as it is not reasonable to measure antigen presentation in every experimental setting.

Our collaborative research has already demonstrated the potential of OSUMMER cells to optimize treatment strategies for NRAS mutant melanomas. For example, using UV-accelerated *TN^61R^* tumor allografts, Echevarria-Vargas et al. showed the synergistic effect of co-targeting Bromodomain and Extra-terminal Domain (BET) and the Mitogen-activated Protein Kinase (MAPK) pathway in NRAS-mutant melanoma (Echevarria-Vargas et al., 2018). Furthermore, the OSUMMER.12 and 13 lines were used to show prolonged tumor responses when targeted therapy was given after anti-PD1—a finding now backed by clinical evidence (Atkins et al., 2022; Phadke et al., 2021). With broad dissemination of the OSUMMER cell lines, we hope to fill a major gap in the field and learn more about immunological profiles and predictors of response to immune, targeted, and combination therapies in NRAS-mutant melanoma.

---

**Supplementary Figure S1:**
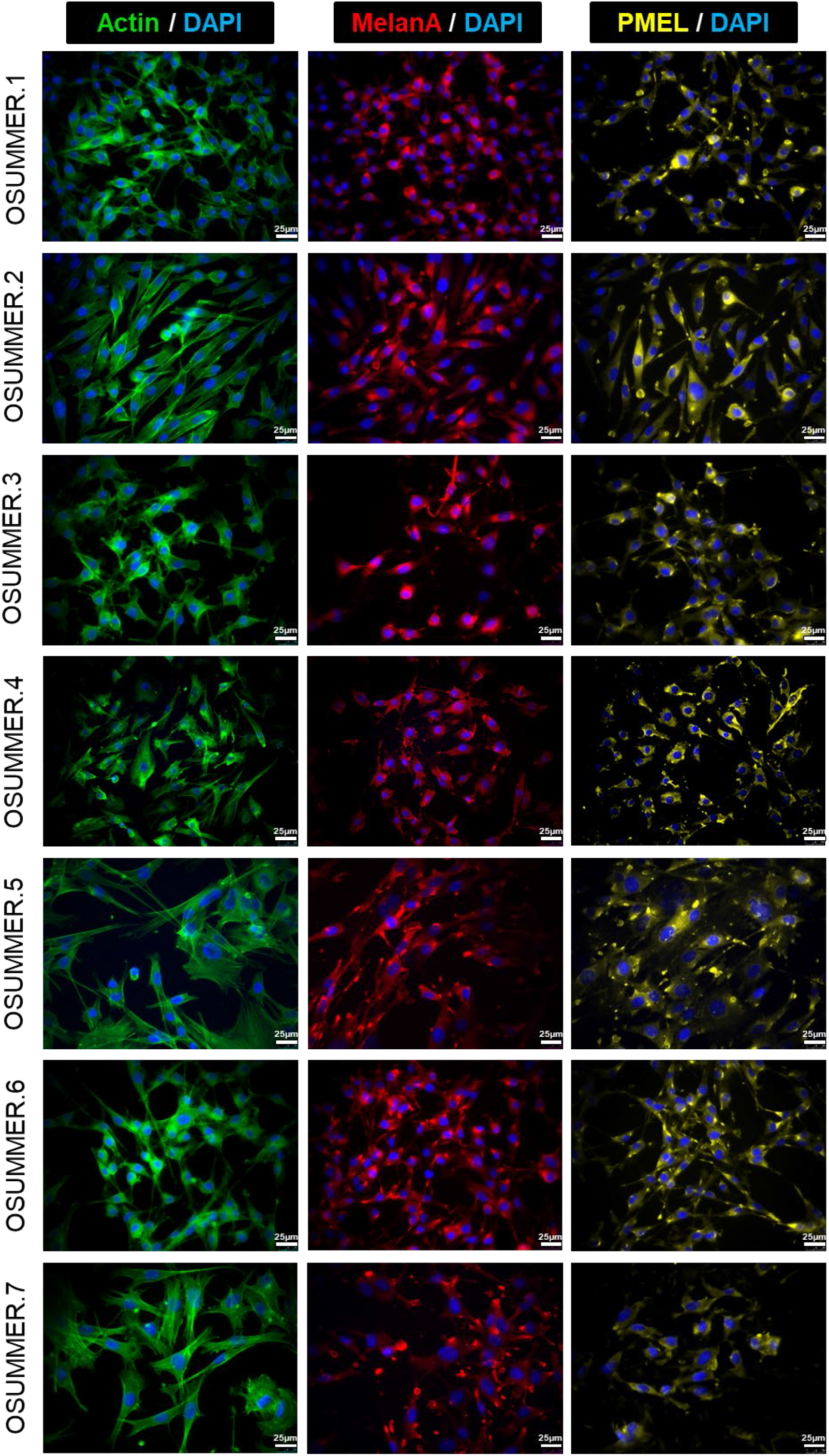
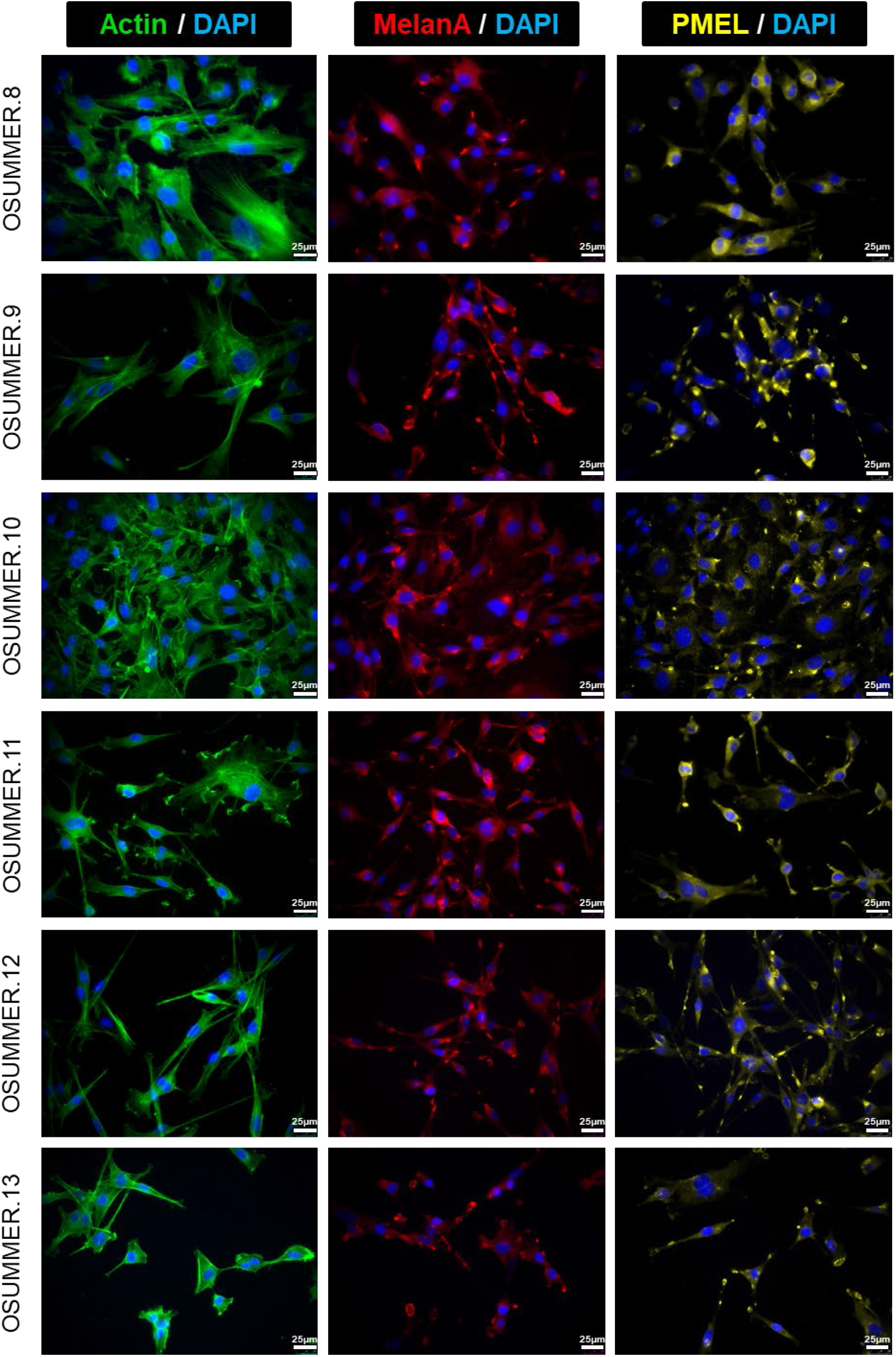
OSUMMER cell lines express the melanocytic markers, MelanA and PMEL. Shown are 40x images of each OSUMMER cell line cultured in vitro and then stained for Actin (green), MelanA (red), PMEL (yellow), and DAPI (blue). Scale bars represent 25 μm.

**Supplemental Figure S2.**
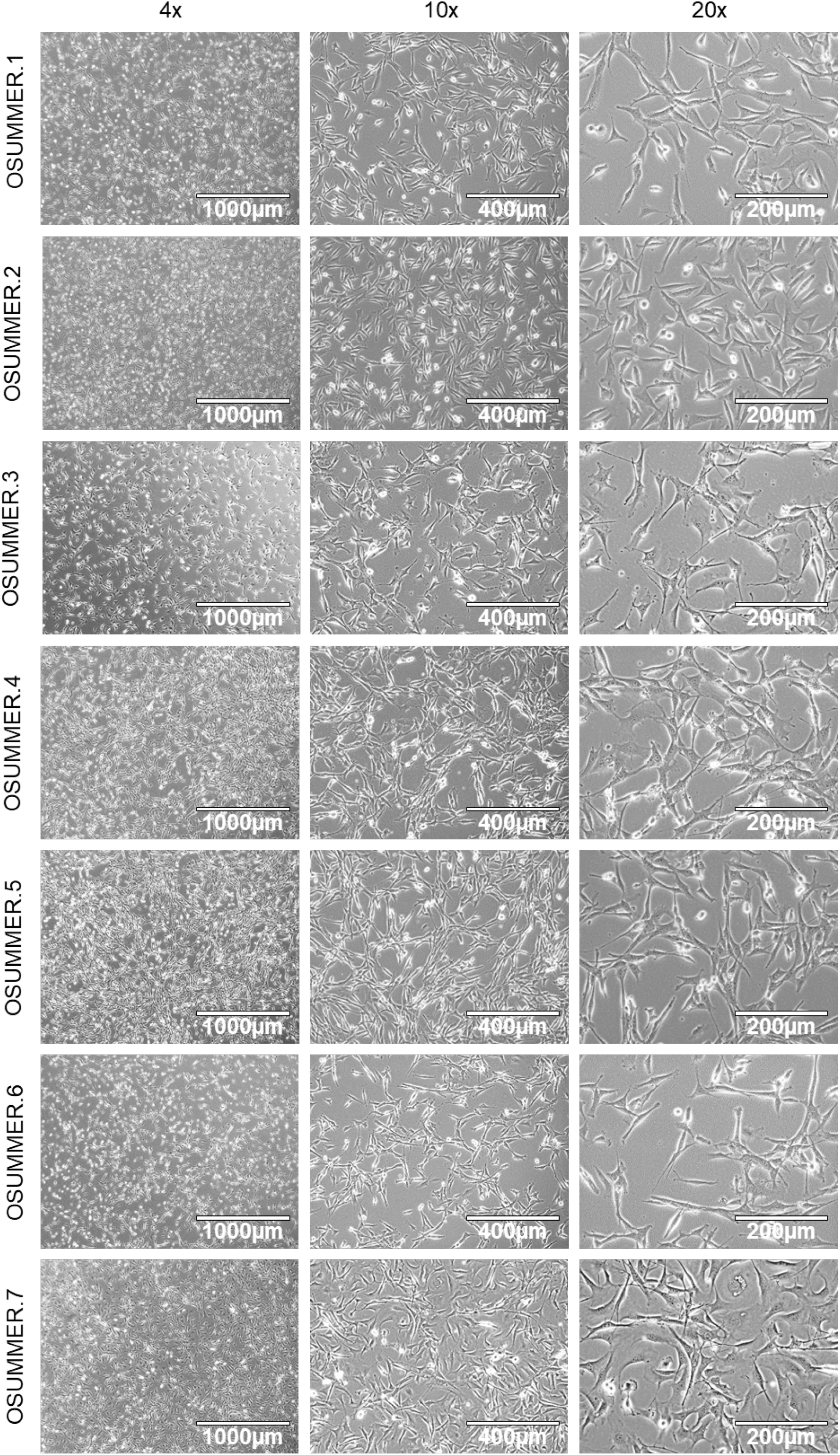
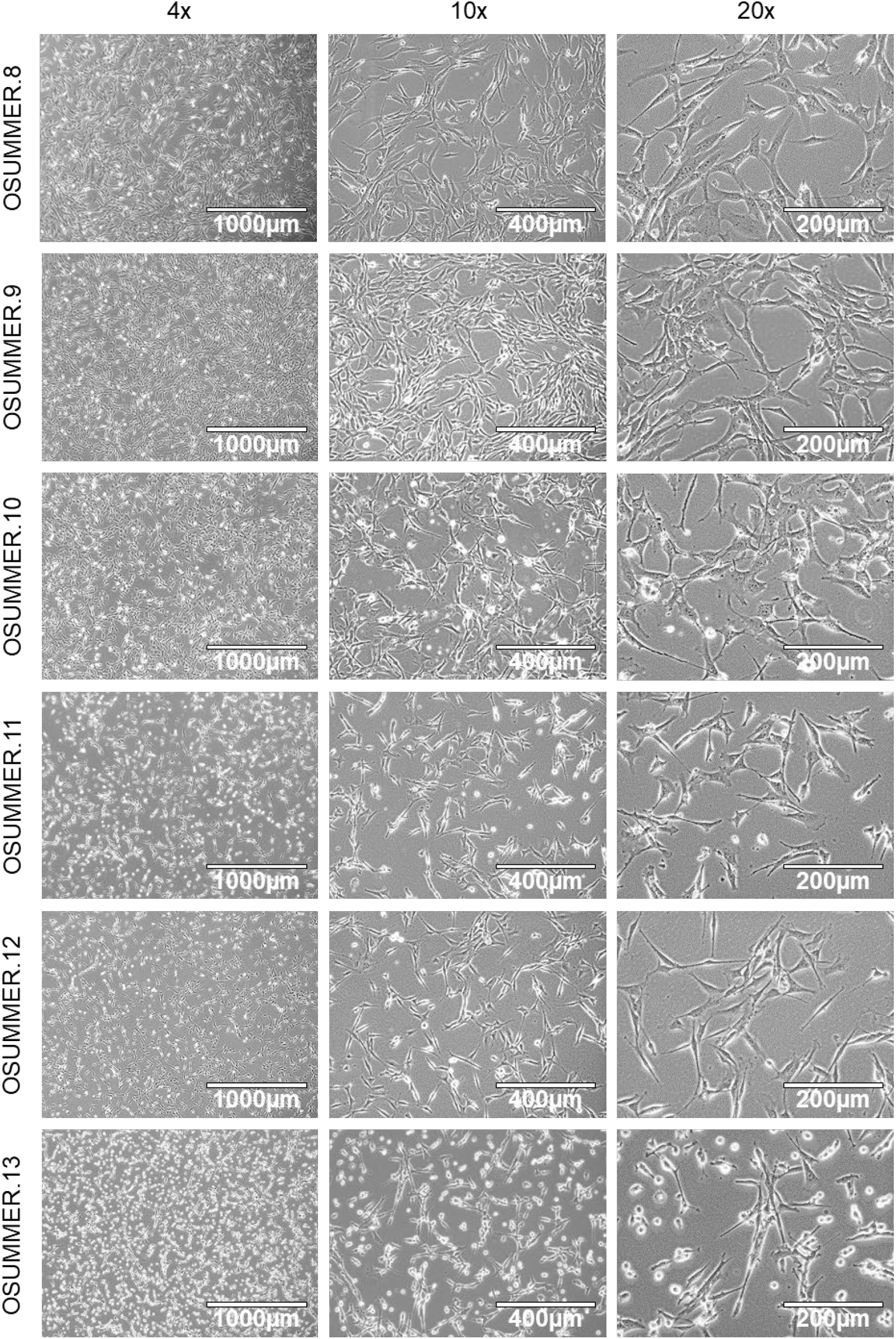
Representative brightfield images of OSUMMER cell lines grown in vitro. Brightfield images of OSUMMER cell lines cultured in vitro. Images were taken at 4x, 10x, and 20x magnification with scale bars representing 1000, 400 or 200 μm as indicated.

**Supplementary Figure S3.**
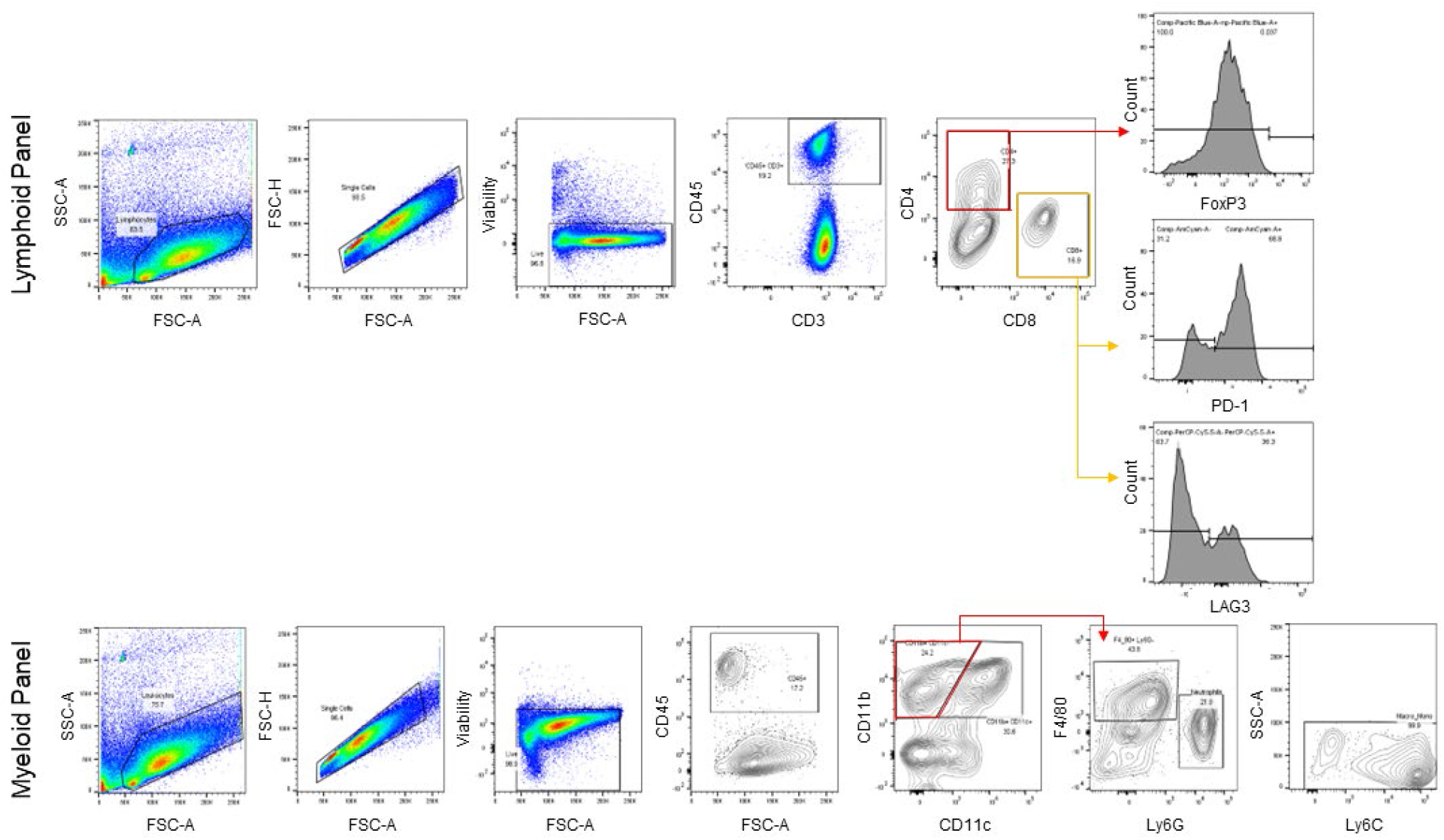
Representative flow cytometry gating schemes for the myeloid and lymphoid panels.

